# Flight demand and environmental niche are associated with molecular evolutionary rates in a large avian radiation

**DOI:** 10.1101/2020.01.16.908368

**Authors:** David A. Duchene, Paola Montoya, Santiago Claramunt, Daniel A. Cadena

## Abstract

Among the macroevolutionary drivers of molecular evolutionary rates, metabolic demands and environmental energy have been a central topic of discussion. The large number of studies examining these associations have found mixed results, and have rarely explored the interactions among various factors impacting molecular evolutionary rates. Taking the diverse avian family Furnariidae as a case study, we examined the association between several estimates of molecular evolutionary rates with proxies of metabolic demands imposed by flight (wing loading and the hand-wing index) and proxies of environmental energy across the geographic ranges of species (temperature and UV radiation). We found evidence that species that fly less have greater wing loading and this is associated with accelerated rates of mutation. An elongated wing morphology is associated with greater flight activity and with molecular signatures of positive selection or reduced population sizes. Meanwhile, environmental temperature and UV radiation interact to explain molecular rates at sites affected by selection and population size, contrary to the expectation of their impact on mutation rates. Our results suggest that the demands of flight and environmental energy pose multiple evolutionary pressures on the genome either by driving mutation rates or via their association with natural selection or population size.

## Introduction

The factors driving molecular evolution are a long standing matter of interest in the biological sciences. Some of the most studied correlates of rates of molecular evolution on a macroevolutionary scale include life-history traits (Bromham 2011), metabolic activity (e.g., Garcia-Porta et al. 2019), and environmental energy (Wright et al. 2006). These factors are also likely to interact, with the evolutionary outcome depending on the balance between the costs of sources of deleterious mutations and the requirements of the particular lifestyle of species (Bromham 2009). Several studies have shown interactions among multiple correlates of molecular evolution (Lanfear et al. 2013; Bromham et al. 2015), but there is still a limited understanding of the relative contribution of biological versus environmental factors to rates of molecular evolution.

Metabolic rate is an important putative driver of molecular evolutionary rates because of the mutagenic effects of metabolism (Gillooly et al. 2005; Allen et al. 2006). Specifically, metabolism produces oxygen and nitrogen free-radicals (Barja 1999) which may cause novel mutations through damage to DNA (Cooke et al. 2003). Mass-specific metabolic rate is inversely associated with body mass (Martin and Palumbi 1993), which provides one explanation for the accelerated rates of molecular evolution in small vertebrates compared to large ones. While there is some evidence that metabolic rates are indeed associated with rates of molecular evolution (Bleiweiss 1998; Gillooly et al. 2005, 2007; Garcia-Porta et al. 2019), multiple studies have not found such an association (Mooers and Harvey 1994; Bromham et al. 1996; Lanfear et al. 2007; Galtier et al. 2009; Rolland et al. 2016).

An explanation for the mixed evidence of the impact of metabolism on molecular evolution is that there has been a heavy focus on body size and temperature as proxies of metabolic rate (Gillooly et al. 2001), yet multiple biological factors can impact metabolic requirements. For example, flight in birds is an extreme form of endurance exercise (Weber 2009) and is associated with high metabolic rates (Butler and Woakes 1990; Suarez 1992; Ward et al. 2001, 2002), high respiratory rates (Guglielmo et al. 2002), high rates of catabolism of lipids and proteins (Jenni-Eiermann et al. 2002), and oxidative stress (Jenni-Eiermann et al. 2014). Therefore, the high exposure to reactive species of oxygen and nitrogen in birds with high flight demands (Jenni-Eiermann et al. 2014) might have a mutagenic effect leading to faster rates of molecular evolution beyond the effects of temperature and body mass.

Alternatively, molecular evolution might not be dominated by mutagens driving the rate of mutation, but by the origin of adaptations to highly demanding lifestyles. Among the many adaptations in avian lineages for sustaining long periods of flight with high efficiency are a high capillary density in flight muscles (Bishop et al. 1995; Maillet and Weber 2007), low wing loading (Norberg 1995; Alerstam et al. 2007), high wing aspect-ratio (Norberg 1990; Pennycuick 2008; Taylor and Thomas 2014), and major changes in skeletal structures and tissues (e.g., West *et al.* 2007; Dumont 2010). Flight demand might also lead to adaptations for lifestyles with high metabolic turnover (i.e., a ‘fast pace of life’): birds that undergo high flight demand are likely adapted to having rapid growth, high fecundity, and short lifespans (Pérez-Tris and Tellería 2002; Ricklefs and Wikelski 2002; Wikelski et al. 2003; but see Winger and Pegan 2020). The signatures for changes in molecular evolutionary rates as driven by flight demand might therefore be predominant in regions of the genome influenced primarily by selection, rather than mutation. In the absence of whole-genome sequences for large numbers of closely related species, one window into genomic regions under different selective regimes is offered by examining the relative contribution of synonymous and non-synonymous substitutions in coding regions (Ohta 1992; Lanfear et al. 2010b).

Another widely studied factor that might drive molecular evolutionary rates is environmental energy through its mutagenic effect on the genome (Martin and Palumbi 1993; Allen et al. 2006). Studies in organisms including plants (Davies et al. 2004; Wright et al. 2006), marine fishes (Wright et al. 2010b), and lizards (Garcia-Porta et al. 2019) have found associations between environmental energy and molecular rates. The link between energy and molecular evolution is often inferred using environmental temperature as a proxy of available energy (Wright et al. 2006). Other studies have focused on UV radiation (e.g., Davies *et al.* 2004), a well understood mutagen (Pawlowski et al. 1997). Temperature and UV can also interact, such that DNA repair is less effective at high UV radiation but low temperatures (MacFadyen et al. 2004). However, it is unclear how UV and its interaction with temperature might impact the germline in many organisms, like endothermic vertebrates (Alton and Franklin 2012; Ghanizadeh Kazerouni et al. 2016).

Environmental energy might also affect molecular evolution by mediating the fate of novel mutations, as opposed to affecting basal mutation rates. For example, temperature might influence the fitness effects of mutations, with maximal fitness occurring at around global maximal temperatures (Puurtinen et al. 2016). Energy might also facilitate the accumulation of biomass and allow high-energy environments to sustain larger numbers of individuals (Wright 1983; Currie 1991; Willig et al. 2003); in larger populations, selection is more efficient and novel beneficial mutations are more likely to reach fixation rapidly (Kimura 1968; Ohta 1992; Lanfear et al. 2014). High energy might also lead to reduced investment in thermoregulation, allowing limited resources to be used in other activities, such as reproduction, and ultimately allowing for novel adaptations (Turner et al. 1988).

We examined the association between the rates of molecular evolution and the metabolic demands imposed by flight and environmental energy across species of the avian family Furnariidae, a large radiation including the Neotropical ovenbirds and woodcreepers (Sibley and Monroe 1990; Remsen Jr et al. 2012). Furnariids have undergone fast and consistent diversification over the last 30My (Derryberry et al. 2011; Harvey et al. 2020), and inhabit a broad range of habitats in South and Central America (Claramunt 2010). The species richness of the family and range of habitats it occupies across broad gradients of altitude, latitude, and environmental conditions make it an uniquely diverse clade in which to test the relative association between molecular evolution and physiology and environmental niche (Fjeldså et al. 2005).

We focus on measurements of the flight apparatus, and environmental temperature and UV radiation as proxies of metabolic demand and environmental energy, respectively. Using data for more than half of the species of Furnariidae (63%) we provide the first near-species-level examination of the correlates of molecular evolution in a large vertebrate radiation. We assessed the association between wing morphology and environmental energy with molecular evolutionary rates in a set of nuclear and mitochondrial genomic regions. Individual tests across synonymous and non-synonymous positions were made to dissect the independent effects on molecular signatures of selection or population size and mutation rate (e.g., Lanfear et al. 2010a; Duchêne and Bromham 2013). The findings suggest a nuanced picture of the impact of flight demands and environmental energy on genome evolution, and highlight several questions that need to be addressed in genome-level studies.

## Methods

### Data collection

We used data on wing shape and body mass obtained from museum specimens as proxies of the flight demands for 290 species in Furnariidae (Claramunt et al. 2012b). Specifically, we measured the distance from the carpal joint to the tip of the longest primary feather (*WL,* the traditional “wing length” measurement) and the distance from the carpal joint to the tip of the first secondary feather (*S1*) and obtained body mass from specimen labels. With these measurements we calculated the hand-wing index as 100(*WL* – *S1*)/*WL* and the wing loading as *M/3 WLS1,* where *M* is the body mass and 3 *WLS1* is an estimate of the total wing area (Claramunt and Wright 2017). The hand-wing index is proportional to the aspect ratio of the wings in furnariids (Claramunt et al. 2012b) and therefore it is related to long-distance flight efficiency (Norberg 1990; Pennycuick 2008). Wing loading is related to the power required for flight regardless of the distance travelled, and is thus associated with the metabolic demand per unit of time in flight (Rayner 1988; Norberg 1990; Bowlin 2007; Bowlin and Wikelski 2008).

All else being equal, wings with high aspect ratio and low wing loading are expected to result in lower metabolic demands. However, across all birds, the relationship between the efficiency of flight and lifetime metabolic demands may be positively correlated because the species that show highly flight-efficient morphologies are also those that fly more frequently and longer distances (Rayner 1988; Norberg 1990; Pennycuick 2008). Meanwhile, species with the least efficient flight morphologies tend to avoid flight and thus experience lower metabolic demands. Therefore, we first examined the relationship between flight efficiency and flight behaviour in Furnariidae to validate the use of flight morphology as a proxy for metabolic demands.

We used basic natural history information to characterize flight behavior of selected species of Furnariidae. For well-understood species in each genus, we collected information about seasonal movement (0 = year-round sedentary or territorial; 1 = non-breeding seasonal displacements or nomadism; 2 = migratory), foraging behaviour (0 = flight rarely used during foraging, short flights <10m seldom needed for crossing habitat gaps or commuting from roosting or nesting sites; 1 = flight used sporadically during foraging, for moving through habitats or for commuting; 2 = flight used regularly for all or nearly all foraging activities), and foraging stratum (0 = ground, low dense vegetation or forest understory; 1 = midstory; 2 = canopy). We used the sum of scores divided by the maximum value (6) as an index of the lifelong flight habits of species, and used phylogenetic regression to examine the association between these data and each of hand-wing index and wing loading (see methods in *Regression analyses*). While these data are useful to verify expectations about wing metrics as proxies of metabolic demand, the flight score was not tested for its association with molecular rates due to its restricted sample size and approximate nature.

We collected data on environmental temperature and UV radiation across the ranges of the species in the family Furnariidae. For all species, we downloaded georeferenced records from GBIF (Global Global Biodiversity Information Facility, https://www.gbif.org/) and VertNet (http://portal.vertnet.org/), discarding duplicated records and those outside the known distribution range for each species according to expert-based maps (del Hoyo et al. 2016). Using 145,216 vetted records (mean per species = ~615), we estimated the mean and standard deviation for the annual mean temperature from WorldClim data (Fick and Hijmans 2017) at 30 arc-sec resolution, and for the annual mean UV-B from gIUV (Beckmann et al. 2014) with 15 arc-min resolution. Among the environmental variables available, temperature and UV provide a broad description of environmental energy, yet they are not simply correlated to each other; these variables follow a triangular-shaped association, where high values of UV can also occur in many habitats with low temperatures (e.g., high altitudes). Because current geographic distributions may differ greatly from distributions in the past, contemporary measurements of environmental variables can introduce noise to subsequent phylogenetic regression analyses. This noise is likely to increase the Type II error rate but it is unlikely to cause a bias that increases Type I error rate, so analyses of this type of data sets are in fact conservative (Davies et al. 2004).

Genomic data were taken from a published phylogenetic study of Furnariidae analysing sequence data from mitochondrial and nuclear markers collected for nearly every species in the family (Derryberry et al. 2011). Using data from two nuclear and three mitochondrial loci (4023 nt of *recombination activating genes 1* and *2;* and 2076 nt of *NADH dehydrogenase subunits 2* and *3,* and *cytochrome oxidase subunit 2,* respectively) from the source phylogenetic study of Furnariidae, we identified species for which data were available for environmental variables, wing loading, and hand-wing index. Our analyses focused exclusively on species with measurements of wing metrics from known museum specimens, and species for which molecular rates could be estimated with confidence (non-zero values, see below; *N* nuclear data = 55; *N* mitochondrial data = 184; Supplementary Figure S1). A nuclear intron used in the original study was not included. The molecular data and variables used in subsequent regression analyses are available online (github.com/duchene/furnariidae_rates).

### Estimates of rates of molecular evolution

Absolute rates of expected synonymous and non-synonymous substitutions per nucleotide site per million years were estimated form the sequence alignments along the branches of an existing time-calibrated tree of Furnariidae (Derryberry et al. 2011). To make reliable analyses of molecular evolutionary rates, we first tested for potential biases from the molecular substitution model using simulation-based tests of model adequacy and substitution saturation as implemented in the software PhyloMAd (Duchêne et al. 2018). This procedure assesses whether empirical data adhere to the assumptions made by the model by comparing them with data simulated under the model. We then estimated the expected number of synonymous and non-synonymous substitutions along branches (expressed as branch lengths) using the MG94 model of codon evolution with codon frequencies estimated from the data (Muse and Gaut 1994), implemented in HyPhy v2.5 (Pond et al. 2005). This model was used in combination with the best-fitting nucleotide substitution model of the GTR+Γ family (Tavaré 1986). Synonymous and non-synonymous substitutions along branches were estimated independently for mitochondrial and nuclear data. Terminal molecular branch-length estimates were divided by the published time-estimate for each branch (Derryberry et al. 2011) and taken as the mean molecular evolutionary rates of species branches.

The resulting synonymous substitution rates (*d*_S_) are the rates of molecular changes not influencing the amino acid being coded, and are proportional to the mutation rate under the condition of no bias in codon usage (Kimura 1968). Meanwhile, non-synonymous substitution rates (*d*_N_) represent rates of amino-acid substitutions, and thus reflect the mutation rate and the interaction between selection and population size (Ohta 1992). The *d*_N_/*d*_S_ ratio is therefore expected to reflect the effects from selection and population size, excluding the effect of the mutation rate.

### Regression analyses

We used phylogenetic generalized least squares (PGLS) regression models to examine the relationship between wing loading, the hand-wing index and variables describing environmental energy, and molecular evolutionary rates. Hypotheses were tested independently using nuclear and mitochondrial data, and using each of *d*_N_, *d*_S_ and *d*_N_/*d*_S_ rates as response variables. The explanatory variables in the model included the hand-wing index, mean environmental temperature, and mean UV radiation. We also included body mass as an explanatory variable in the models containing the hand-wing index, because body mass affects most life-history traits (e.g., generation time, lifespan) and demography, so it has a possible confounding effect of the association between the hand-wing index, environmental variables, and molecular evolution (Bromham 2009). Interaction terms included that between body mass and hand-wing index, allowing us to consider differences in biology between species of different body sizes, and the interaction between the environmental temperature and UV radiation. Body mass was not included in models together with wing loading given the strong association between the two variables.

To account for non-independence of data due to relatedness among taxa, we used a species-level phylogenetic estimate extracted from the original phylogenetic study (Derryberry et al. 2011). We normalized the molecular variables using the *normalize* function of the R package *BBmisc* (Bischl et al. 2017), and then performed a Box-Cox transformation on all molecular variables and log transformation on body mass to adhere to least-squares model assumptions. The lambda parameter for the Box-Cox transformation was estimated with *boxcox* function of the *MASS* R package (Venables and Ripley 2002), adding as a constant value the minimum in each variable plus 0.01 on the respective variable, to avoid negative values. PGLS regression was performed under a lambda model of trait evolution, such that we estimated the best fitting value of phylogenetic inertia in the residuals simultaneously with other parameters (Freckleton et al. 2002). The parameters of PGLS regression models were optimized using the *pgls* function of the *caper* R package (Orme et al. 2018), and residuals for each model were assessed visually for normality. Because regression models containing each of the molecular variables and environmental variables were ran twice, we verified that any results remained qualitatively identical by adjusting *p*-values using false discovery rates.

## Results

The use of flight in Furnariidae was negatively associated with wing loading (*p* = 0.002, *R*^2^ = 0.24) and positively associated with the hand-wing index (*p* < 0.001, *R*^2^ = 0.58; Supplementary Figure S2; full results available online, github.com/duchene/furnariidae_rates). Therefore, flight-intensive habits tend to occur in species with low wing loading and high values of the hand-wing index.

Regression analyses revealed a positive association between wing loading and both mitochondrial *d*_N_ (*p* = 0.04, *R*^2^ = 0.05) and *d*_S_ rates (*p* = 0.013, *R*^2^ = 0.05; Table 1; full models results available online, github.com/duchene/furnariidae_rates), indicating faster mutation rates in species with higher wing loadings. We also found a significant positive interaction between the hand-wing index and body mass in explaining mitochondrial molecular evolutionary rates (*d*_N_/*d*_S_, *p* = 0.031, *R*^2^ = 0.04; and *d*_N_, *p* = 0.035, *R*^2^ = 0.06; Table 2), but the interaction was not observed in the proxies of molecular evolution most closely associated with mutation rates (*d*_S_). More specifically, there was a positive association between the hand-wing index and molecular rates in small-bodied species of Furnariidae but a positive association between the hand-wing index and molecular rates in large-bodied species.

**Table 1.** Parameter estimates of the PGLS regression for models testing wing loading and environmental variables as explanatory of molecular rates. A Box-Cox transformation was used for all molecular rates variables.

| Molecular rate response regression terms | Wing loading | Mean annual temperature | Mean UV radiation | Mean Annual Temp * Mean UV radiation | $R^2$ |
| --- | --- | --- | --- | --- | --- |
| Mitochondrial $d_N/d_S$ | 0.02 | 0.027 | -0.073 | -0.084 | 0.011 |
| Mitochondrial $d_N$ | 0.1* | -0.013 | -0.185 | -0.162* | 0.048 |
| Mitochondrial $d_S$ | 0.121* | -0.055 | -0.133 | -0.105 | 0.048 |
| Nuclear $d_N/d_S$ | 0.022 | 0.008 | 0.048 | 0.088 | 0.02 |
| Nuclear $d_N$ | -0.083 | 0.033 | -0.036 | -0.018 | 0.008 |
| Nuclear $d_S$ | -0.171 | -0.025 | -0.031 | -0.092 | 0.055 |

$P$ -values denoted $\cdot \leq 0.1$ , $* \leq 0.05$

**Table 2.**
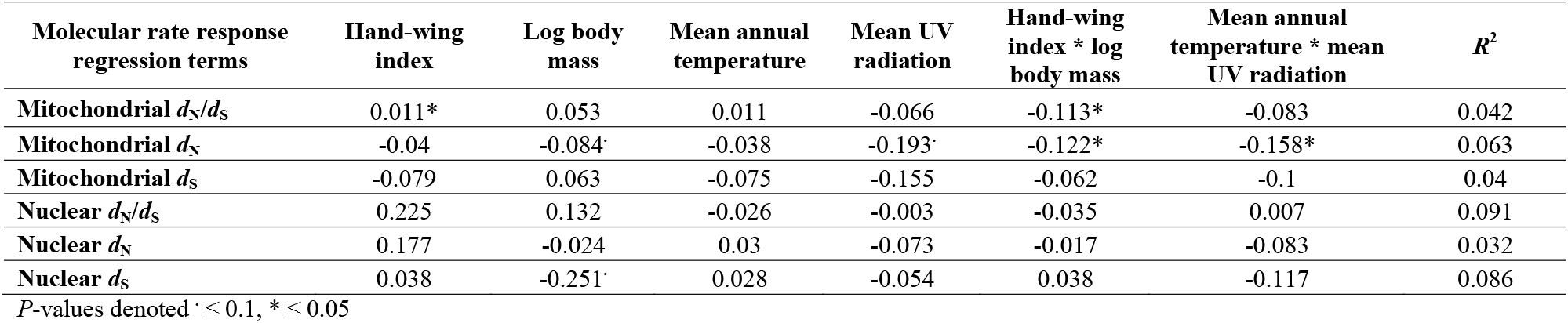
Parameter estimates of the PGLS regression for models testing the hand-wing index and environmental variables as explanatory of molecular rates. A log transformation was used for the mitochondrial *d*_N_/*d*_S_ rates, and a Box-Cox transformation was used for all other molecular rates variables.

| Molecular rate response regression terms | Hand-wing index | Log body mass | Mean annual temperature | Mean UV radiation | Hand-wing index * log body mass | Mean annual temperature * mean UV radiation | $R^2$ |
| --- | --- | --- | --- | --- | --- | --- | --- |
| Mitochondrial $d_N/d_S$ | 0.011* | 0.053 | 0.011 | -0.066 | -0.113* | -0.083 | 0.042 |
| Mitochondrial $d_N$ | -0.04 | -0.084 | -0.038 | -0.193 | -0.122* | -0.158* | 0.063 |
| Mitochondrial $d_S$ | -0.079 | 0.063 | -0.075 | -0.155 | -0.062 | -0.1 | 0.04 |
| Nuclear $d_N/d_S$ | 0.225 | 0.132 | -0.026 | -0.003 | -0.035 | 0.007 | 0.091 |
| Nuclear $d_N$ | 0.177 | -0.024 | 0.03 | -0.073 | -0.017 | -0.083 | 0.032 |
| Nuclear $d_S$ | 0.038 | -0.251 | 0.028 | -0.054 | 0.038 | -0.117 | 0.086 |

$P$ -values denoted $\cdot \leq 0.1$ , $* \leq 0.05$

There was a negative interaction between the two environmental variables in explaining mitochondrial *d*_N_ molecular rates (*p* = 0.033). The direction of this interaction suggests that species of Furnariidae have faster rates of *d*_N_ when environments in their geographic range have relatively low temperatures but high exposure to UV radiation. As is common in analyses of molecular evolutionary rate estimates, we found all linear models to explain low portions of the variation in the data, with *R*^2^ values consistently below 0.1 (Welch and Waxman 2008).

**Figure 1.**
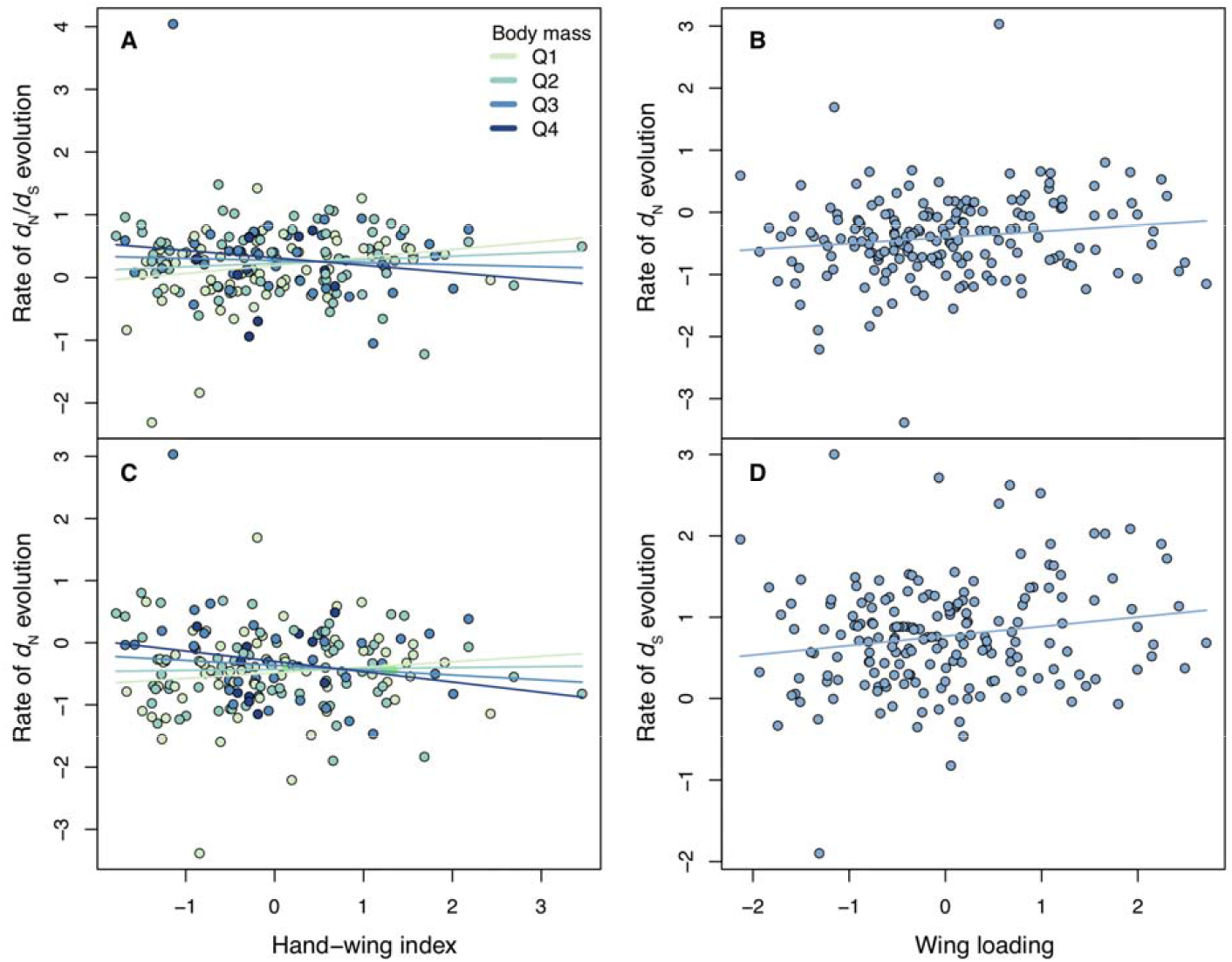
Association between the (A, C) hand-wing index and mitochondrial molecular rates associated with selection or population size; and between (B, D) wing loading and mitochondrial molecular rates associated with mutation rates across species of the avian family Furnariidae. Colours (A, C) represent mid-quartile values of body mass, where darker colours indicate species with greater body mass. The predicted regression lines are also coloured for each of the mid-quartile values of body mass. Molecular rates are shown normalized and under a Box-Cox transformation.

**Figure 2.**
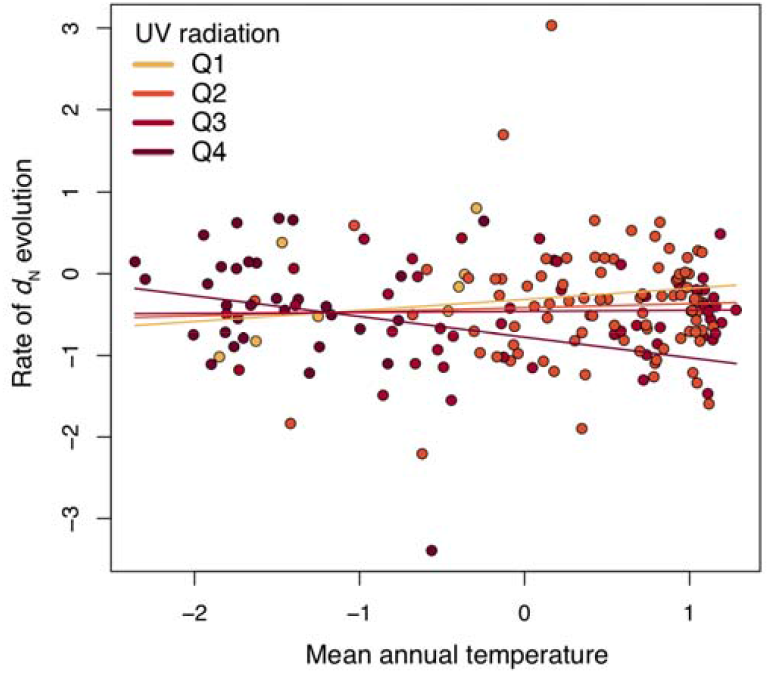
Association between environmental variables and mitochondrial *d*_N_ rates across species of the avian family Furnariidae. Colours represent mid-quartile values of UV radiation, where darker colours indicate greater values. Molecular *d*_N_ rates are shown normalized and under a Box-Cox transformation.

## Discussion

Our study uncovered associations among wing metrics as proxies of the metabolic demands related to flight, environmental temperature and UV radiation with molecular evolutionary rates across the Neotropical avian family Furnariidae. We found evidence of significantly higher mutation rates in species with higher wing loading, despite being associated with infrequent flight habits. This result is consistent with the idea that highly demanding bursts of metabolic activity, even if infrequent, can increase germline mutation rate. Our analyses also reveal a positive association between wing shape as measured by the hand-wing index and non-synonymous substitution rates, primarily in small-bodied species. The effects of flight demands as indexed by wing shape and environmental energy on the genome might therefore occur via selective pressures or effects on population size. This result is consistent with previous studies showing that flight has led to a broad range of genomic as well as physiological adaptations (Tobalske et al. 2003; Wright et al. 2014), and with evidence that metabolic rates do not have a simple association with molecular rates (Lanfear et al. 2007). Our data also offer support for an association between environmental energy and molecular evolutionary rates in birds (Gillman et al. 2012), and for a negative interaction between temperature and UV radiation, in agreement with previous evidence that these two measures of environmental energy can compensate each other (Hoffman et al. 2003; Jin et al. 2019). Future genome-wide studies will be instrumental to describe molecular regions most impacted by the mutation pressures and selection or population size effects arising from energetically demanding flight and high-energy environments (Zhang et al. 2014).

Theory predicts that metabolic rates can drive basal mutation rates (Allen et al. 2002; Gillooly et al. 2005), and this is supported by our findings of faster mutation rates in species with greater wing loading. However, the opposite might also be true, whereby factors associated with low wing loading place greater pressure to reduce mutation rates. Most organisms likely exist at an upper limit of mutations, such that even a minor increase in mutation rate may lead to an unsustainable loss of fitness (Bromham 2011). For example, large-bodied mammals might accumulate more mutations than small-bodied mammals because their long lifespan allows for more generations of cells producing gametes, and this might interact with the smaller population sizes of larger organisms to result in faster accumulation of deleterious changes (Bromham 2009). Nonetheless, instead of having a fast rate of accumulation of changes, large-bodied mammals have relatively slow molecular evolutionary rates, possibly reflecting high fitness costs of deleterious changes (Lynch 2010). Fast metabolic rates in birds with intense flights might pose a similarly increased selective pressure to avoid deleterious mutations, reducing the basal mutation rate. The positive association between wing loading and a signal of mutation rates might therefore be caused by an increased exposure to mutagens from bursts of metabolism in taxa with high wing loading (e.g., Lanfear *et al.* 2007; Rolland *et al.* 2016), or a pressure in taxa with low wing loading (and frequent flight habits) to reduce mutation rates.

The differences in the results found using wing loading and hand-wing index can be explained by the differences between the two metrics, given they capture different dimensions of flight habits. Our data reveal that the two metrics follow opposite associations with flight habits. Species with high wing loading also tend to have greater body masses and are generally more sedentary. Meanwhile, species with high hand-wing indices fly more frequently and also more economically compared with species with low hand-wing indices. Accordingly, wing loading is more likely to correlate with the immediate metabolic demand caused by bursts of flight, whereas the hand-wing index is more likely to correlate with the demands of regular or long-duration flight. Experimental tests of these findings would be a fruitful area of future research (e.g., Reynolds et al. 2014).

Instead of finding effects suggesting that measures of environmental energy are directly associated with the rate of mutation, we found evidence that temperature and UV radiation can interact to accelerate evolutionary rates in gene regions that can change the coded amino-acids. Previous research has found this interaction to impact mutation rates: low temperatures have a negative impact on DNA repair mechanisms and interact with the mutagenic impact of high exposure to UV radiation (MacFadyen et al. 2004). Our finding of an association between flight demand and evolution at regions associated with selection and population size suggests that the interaction influences molecular evolution through mechanisms other than DNA repair. This might be explained by the ecological challenges posed by environments with low temperatures and high UV radiation, such as the reduced oxygen pressure at high altitudes. If this the case, then habitats at extremely high or low energy levels might either drive novel adaptations or maintain a reduced population size, elevating non-synonymous substitution rates.

Independent sources of evidence show that metabolism has placed a set of selection pressures on bird physiology and genomics. For instance, powered flight requires high-energy output (Suarez 1992) and metabolic efficiency (Kvist et al. 2001; Morris et al. 2010), and selection related to the demands of flight is thought to influence genome evolution (Hughes and Hughes 1995; Kapusta et al. 2017). A reduction in genome size is likely to have preceded the emergence of flight in vertebrates (Organ and Shedlock 2009), yet further reductions are thought to be associated with adaptations for flight, including rapid gene regulation (Organ et al. 2007; Zhang and Edwards 2012) and the demands of particular flight modes (Wright et al. 2014). This broad range of evolutionary changes in the genome offers an explanation for our results of increased rates in genomic regions associated with selective pressures in the mitochondria. Instead of placing pressure on the genome to remain unchanged, the wide-ranging adaptations associated with flight demands have likely spurred mitochondrial evolution. Such adaptations include changes in physiology and behavior associated with flight and with the remarkable opportunities for niche differentiation that flying promotes (e.g., Gavrilov 2011; Claramunt *et al.* 2012a; Sol *et al.* 2012). Flight is likely to be a major driver of evolutionary change across multiple genes (Zhang et al. 2014), and exploring larger nuclear regions is likely to provide further insight onto the relative impacts of flight and the environment on avian genome evolution.

Selection and population size often interact in complex ways to influence substitution rates (Lanfear et al. 2014). For example, the relationship between population size and substitution rate takes drastically different shapes depending on whether a genomic region is under positive or negative selection. In the case of regions largely undergoing negative selection (e.g., many protein-coding loci), large population sizes have a greater efficiency of selection, resulting in relatively low substitution rates even under a constant selection coefficient. This means that our data cannot determine whether the hand wing index is associated with substitution rates via an effect on selection or on population size. For instance, our results may reflect a negative association between population size and both the hand-wing index and some measures of environmental energy. The furnariid miners (*Geositta*) and palm creepers (*Berlepschia*) are examples of taxa with a high hand-wing index that are also likely to have low population density. Similarly, tropical regions have previously been associated with lower population density compared with regions with lower environmental energy (Lawrence and Fraser 2020). Some alternatives that might disentangle effects of selection and population size on molecular evolution in future research include the incorporation of independent estimates of population size, or an extensive exploration of the relative impacts of the two factors on molecular evolutionary rates.

## Conclusions

Our study establishes an association between wing morphology and flight habits, and provides evidence that flight and environmental energy can impact genome evolution via an influence on basal mutation rates, as well as via natural selection or effective population size. The metric of flight demand plays an important role in this result. Short but metabolically demanding bursts of flight are associated with fast mutation rates, whereas frequent flights are associated with natural selection or effective population size fluctuations. These results cement the idea that flight demand is associated with a range of adaptations leading to positive selection in large swathes of the genome (Zhang et al. 2014). Meanwhile, our data show further evidence of interacting effects of sources of environmental energy on molecular evolution (e.g., MacFadyen et al. 2004), but likely tied with a range of adaptations or population constraints found at extreme habitats and lifestyles rather than impacting mechanisms of DNA repair. Data from whole genomes and detailed physiology across bird taxa (e.g., Jarvis et al. 2014; Feng et al. 2020; Sheard et al. 2020) will bring a more complete picture of the impact of metabolism, population size, and the environment on avian genomic evolution.

## Supporting information

**Figure S1.**
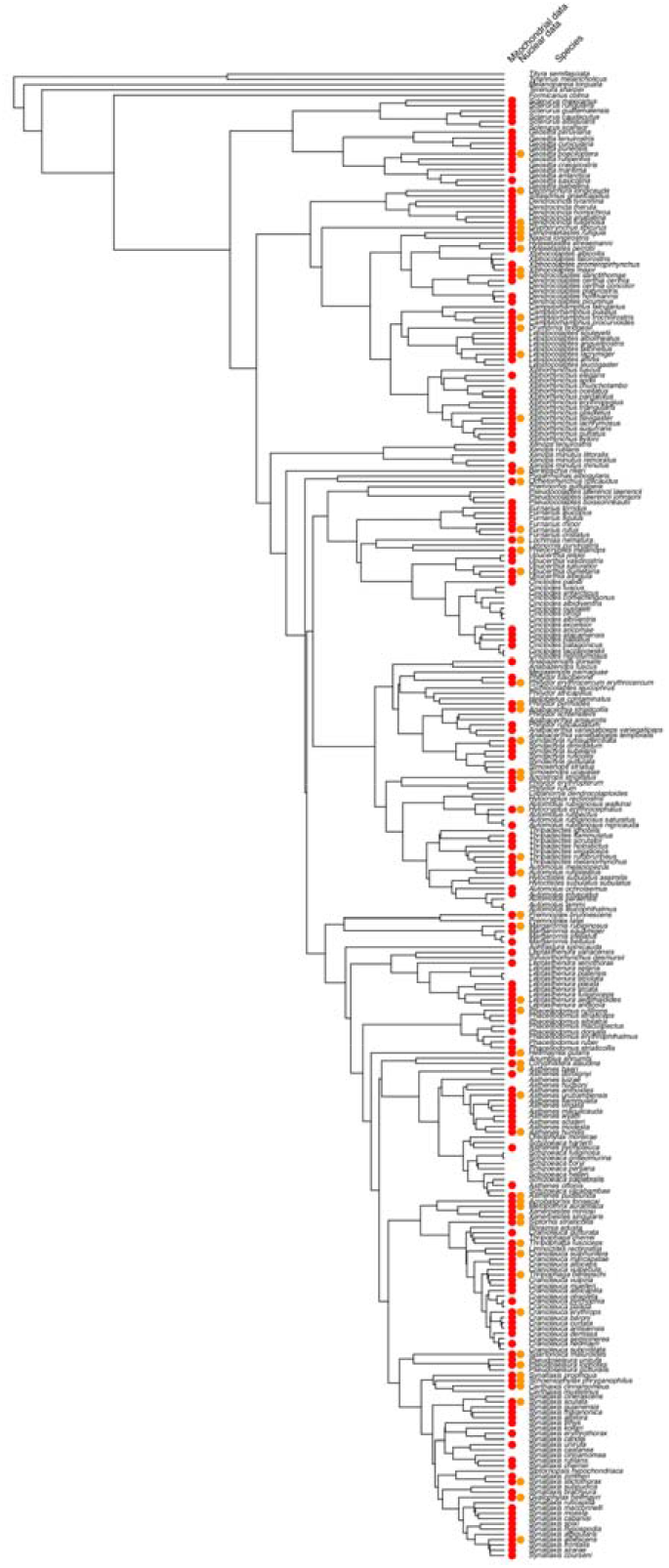
Phylogenetic estimate of the family Furnariidae (Derryberry et al. 2011), showing the taxa included in regression models with mitochondrial (red) and nuclear (orange) data. Inclusion depended on availability of data on all of the variables describing environment, wing loading, hand-wing index, and molecular evolutionary rates.

**Figure S2.**
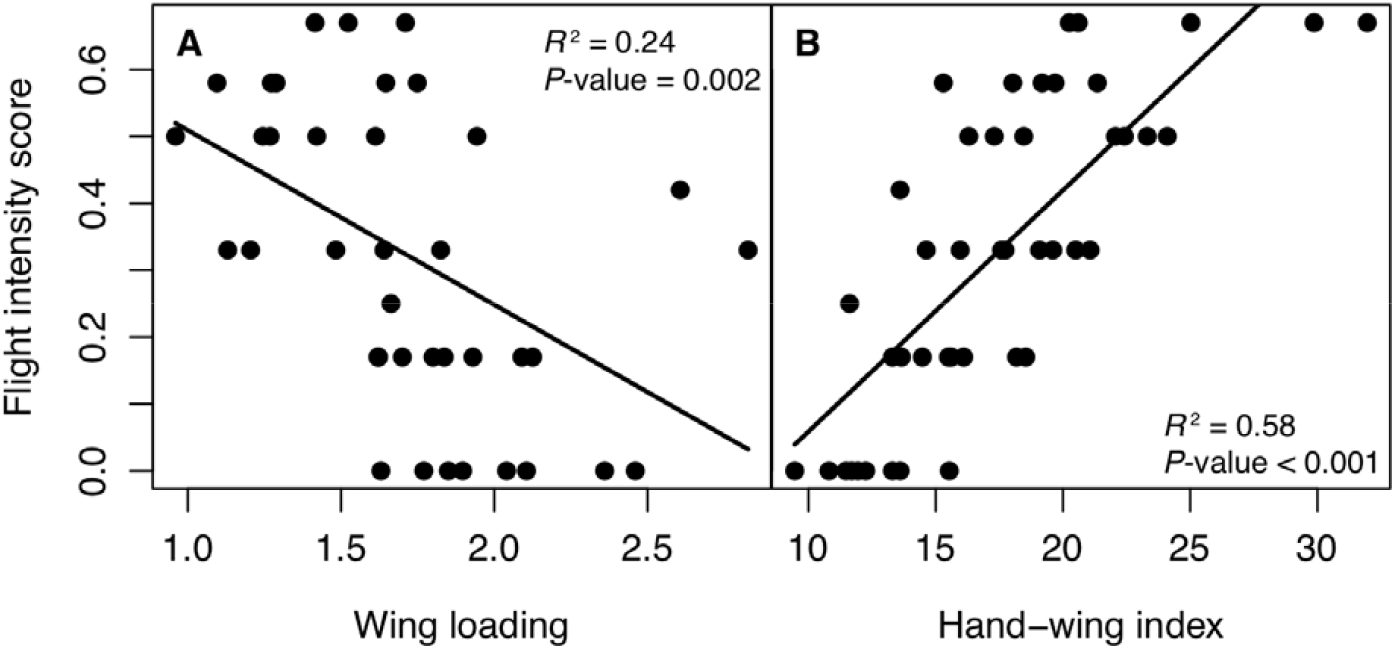
Association between an approximate score of flight-intensive habits (0 = limited flight habits, 1 = intensive flight habits; see Methods section) with (A) wing loading and (B) hand-wing index across 44 genera of the avian family Furnariidae. Lines show the fitted PGLS regression model.

